# Deep6: Classification of Metatranscriptomic Sequences into Cellular Empires and Viral Realms Using Deep Learning Models

**DOI:** 10.1101/2022.09.13.507819

**Authors:** Jan F. Finke, Colleen Kellogg, Curtis A. Suttle

**Affiliations:** Hakai Institute, PO Box 309, Heriot Bay, BC, V0P1H0, Canada; Department of Earth, Ocean and Atmospheric Sciences, The University of British Columbia, Vancouver, BC, V6T1Z4, Canada; Department of Microbiology and Immunology, The University of British Columbia, Vancouver, BC V6T1Z3, Canada; Department of Botany, The University of British Columbia, Vancouver, BC V6T1Z4, Canada; Institute for the Oceans and Fisheries, The University of British Columbia, Vancouver, BC, V6T1Z4, Canada

**Author notes:** Corresponding Authors (Jan F. Finke) and (Curtis A. Suttle).

## Abstract

Metatranscriptomic data is increasingly used to study viral diversity and activity; yet, identifying and taxonomically assigning viral sequences is still challenging. Deep6 is a deep-learning model that classifies metatranscriptomic sequences into six groups: prokaryotes, eukaryotes, or one of the four viral realms. Deep6 is trained on reference coding sequences, but classification of query sequences is done reference-independent and alignment-free. The provided model is optimized for marine samples and can process sequences as short as 250 nucleotides. Average accuracies range from 0.87 to 0.97 depending on sequence length. Additionally, Deep6 includes scripts to easily encode and train custom models for other environments.

---

Marine viruses are agents of microbial mortality, drive nutrient cycles and represent a vast genetic diversity (Suttle 2005). However, viral reference databases are incomplete and studying viral diversity and activity remains challenging. Metatranscriptomes are increasingly used to simultaneously profile biological functions and taxonomic information of cellular and viral origin in the same samples, providing a comprehensive overview (e.g Sutherland et al. 2022). Yet, using current alignment-based methods it is still difficult to identify viral sequences in metatranscriptomic data. Recent machine learning methods such as DeepVirFinder (Ren et al. 2020), DeepMicrobeFinder (Hou et al. 2021) or VirSorter2 (Guo et al. 2021) are designed to compensate for the incomplete viral reference databases. However, DeepVirFinder only discerns between viral vs. non-viral sequences and is trained for viruses that infect prokaryotes. DeepMicrobeFinder discerns sequences among prokaryote and eukaryote hosts, their viruses, and plasmids, but does not specify the viral realm of sequences. VirSorter2 does discern sequences among non-viral and different viral groups, but still relies on reference databases and struggles with sequences shorter than 3000 nt. Here we present Deep6, a reference-independent and alignment-free deep-learning model to predict sequences for all viral realms, optimized for marine samples and short-sequence metatranscriptomic data.

Similar to DeepVirFinder and DeepMicrobeFinder, Deep6 is a deep learning model, but expands on their functionality by classifying sequences into six groups: prokaryotes, eukaryotes, or one of the four viral realms, *Duplodnaviria, Varidnaviria, Monodnaviria* and *Riboviria*. The assumption is that the separation into these six groups reflects inherent differences in nucleotide k-mer patterns among these groups; the model identifies and learns those characteristic patterns from the training sequence data for each class. Deep6 is a multi-class Convolutional Neural Network (CNN) model, consisting of 500 convolutions, 500 dense layers, a default kernel size of ten and a maximum of 40 epochs of training. Four different models accommodate input sequences of 250-499 nt, 500-999 nt, 1000-1499 nt, and >1500 nt to optimize model performance for shorter sequences and to match common distributions of metatranscriptomic sequence lengths. Sequence prediction is done independently of reference databases and can process sequences as short as 250 nt. Cellular training data is compiled from reference Coding Sequences (CDS) for marine eukaryote and prokaryote genera and species based on the Tara Oceans data (Sunagawa et al. 2015). Viral training data is compiled from CDS for all available reference genomes and their neighbour nucleotides (NCBI) for viral orders in the four viral realms. In the training data set each group is represented by between 50,000 and 130,000 randomly selected CDS from the pool of reference data. Additionally to the provided default models, which are currently optimized for marine host virus systems, Deep6 can be readily retrained for custom models for other environments or communities.

The Deep6 repository includes the default models and the primary sequence prediction script. Additionally, the repository includes scripts for batch-encoding datasets and training custom models. Input sequences for prediction or training can be read in FASTA format, and predictions are saved as tab delimited files. The primary prediction script selects the appropriate model for each input sequence length and automatically encodes and predicts the sequence. For each sequence group scores are calculated and then used to derive the final prediction in the downstream analysis. To prepare datasets to train custom models, the batch-encoding script splits sets of CDS into 90% training data and 10% validation data, chunks of CDS of appropriate length for the model in overlapping forward and reverse order are then one-hot encoded. The custom training script feeds training and validation data into the CNN and saves the best model. Models are optimized for accuracy; model performance is assessed by training and validation area under receiver operating characteristic curve (AUROC), average accuracy, and group precision, recall and derived F1-scores. For each final model a confusion matrix of sequence predictions is produced.

The provided default models have training and validation AUROCs ranging from 0.98 to 1.00, and average accuracies ranging from 0.87 to 0.97 (Figure 1). These AUROC values match or exceed those for models for similar sequence lengths in DeepVirFinder and DeepMicrobeFinder. The F1-scores show that model performance varies among groups and with sequence lengths. Overall, models for longer sequences perform better with validation F1-scores between 0.91 and 0.99 for the >1500 nt model and 0.71 and 0.96 for the 250-499 nt model. Across all sequence length models the *Duplodnaviria* is the least predictable group and sequences are primarily miss-predicted as prokaryote. In a cross-validation with 30,000 known CDS with a mean sequence length of 1355 nt 80% of sequences were predicted correctly. Where comparable, F1-scores of 0.79 (eukaryotes), 0.83 (prokaryotes), 0.65 (*Duplodnaviria*), 0.82 (*Varidnaviria*), 0.87 (*Monodnaviria*) and 0.81 (*Riboviria*) match or exceed those reported for 1500 nt log test sequences with VirSorter2. Deep6 represents an effective tool to predict viral sequences from across all realms, optimized for short sequence metatranscriptomic data. Deep6 is a high-throughput program capable of processing about one million sequences per day and can be used to support homology based viral identification and discover novel viruses.

**Figure 1:**
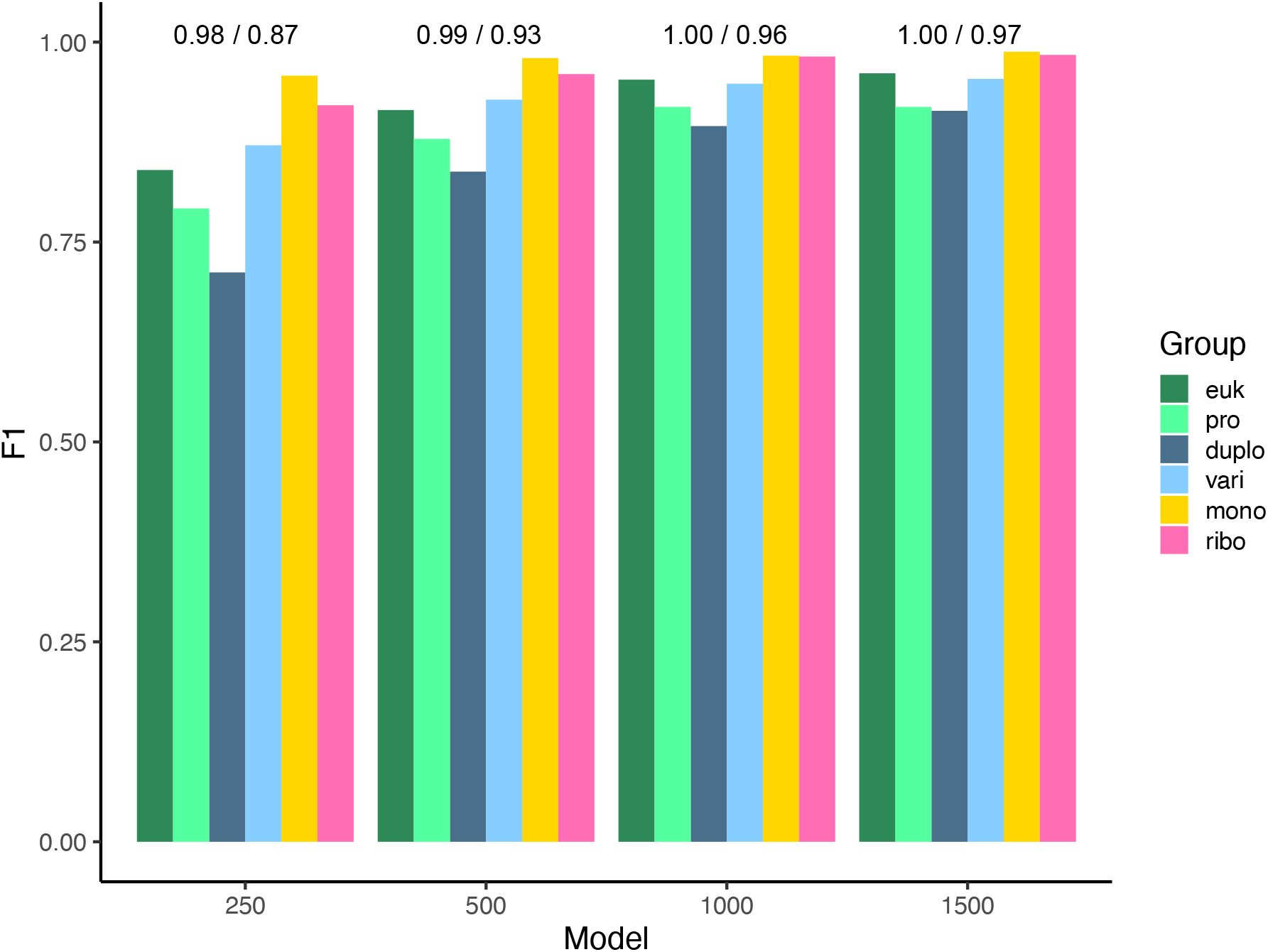
Statistics for the four models and six classification groups. F1-scores are shown for the six classification groups per model. Models are for sequence lengths of 250-499, 500-999, 1000-1499 and >1500 nucleotides. Groups are indicated by colors, eukaryotes (dark green), prokaryotes (light green), *Duplodnaviria* (dark blue), *Varidnaviria* (light blue), *Monodnaviria* (gold) and *Riboviria* (pink). Validation AUROCs and average accuracies per model are shown above bars.

## Availability

Deep6 source code and supporting information is available (under a GNU AGPLv3 license) at: *https://github.com/janfelix/Deep6*. Scripts are written in Python3 and rely on the readily available libraries Biopython and Tensorflow. Environmental files to setup in Anaconda and to install dependencies are also provided. Contact Jan F. Finke for further information on training and crossvalidation datasets.

## Author contributions

JFF, CK and CAS conceived the project and developed the use-case; JFF developed the model and associated code, and compiled the training data; JFF, CK and CAS drafted and edited the manuscript.

## Acknowledgements

JFF was supported during this work by the Tula Foundation. We would like to thank Raymond Ng for advice on model design, and test and training dataset design. We would also like to thank Jonathan Schatz for advice on data sourcing and code development.

## References

Guo, Jiarong, Ben Bolduc, Ahmed A. Zayed, Arvind Varsani, Guillermo Dominguez-Huerta, Tom O. Delmont, Akbar Adjie Pratama, et al. 2021. “VirSorter2: A Multi-Classifier, Expert-Guided Approach to Detect Diverse DNA and RNA Viruses.” Microbiome 9 (1): 1–13. https://doi.org/10.1186/s40168-020-00990-y.

Hou, Shengwei, Siliangyu Cheng, Ting Chen, Jed A. Fuhrman, and Fengzhu Sun. 2021. “DeepMicrobeFinder Sorts Metagenomes into Prokaryotes, Eukaryotes and Viruses, with Marine Applications.” BioRxiv, 1–25. https://www.biorxiv.org/content/10.1101/2021.10.26.466018v1%0A https://www.biorxiv.org/content/10.1101/2021.10.26.466018v1.abstract.

Ren, Jie, Kai Song, Chao Deng, Nathan A. Ahlgren, Jed A. Fuhrman, Yi Li, Xiaohui Xie, Ryan Poplin, and Fengzhu Sun. 2020. “Identifying Viruses from Metagenomic Data by Deep Learning.” Quantitative Biology 8 (1): 64–77.

Sutherland, Ben J.G., Jan F. Finke, Robert Saunders, Snehal Warne, Angela D. Schulze, Jeff H.T. Strohm, Amy M. Chan, Curtis A. Suttle, and Kristina M. Miller. 2022. “Metatranscriptomics reveals a shift in microbial community composition and function during summer months in a coastal marine environment.” Environmental DNA: 1–14. https://doi.org/10.1002/edn3.353

Suttle, Curtis A. 2005. “Viruses in the Sea.” Nature 437 (7057): 356–61. https://doi.org/10.1038/nature04160.

